# Temporal responses of bumblebee gustatory neurons to sugars

**DOI:** 10.1101/2022.02.28.482341

**Authors:** Rachel H. Parkinson, Sébastien C. Kessler, Jennifer Scott, Alexander Simpson, Jennifer Bu, Adam Mahdi, Ashwin Miriyala, Geraldine A. Wright

## Abstract

The sense of taste permits the recognition of valuable nutrients and the avoidance of potential toxins. Models of gustatory coding propose that within modalities (e.g. sweet, bitter, etc.), taste ligands are not distinct stimuli. However, these models are based on data from mice or flies that have omnivorous, non-specialist diets. A specialist feeder might, however, be expected to have acuity within modality if stimulus resolution was critical to survival. Previously, we found that bumblebees have a specialized mechanism for sensing sugars whereby two gustatory receptor neurons (GRNs) within the galeal sensilla of the bees’ mouthparts exhibit burst of spikes. Here, we show that the temporal firing patterns of these GRNs separate sugars into four distinct groups that correlate with sugar nutritional value and palatability. We also identified a third GRN that responded to stimulation with relatively high concentrations of fructose, sucrose, and maltose. Sugars that were non-metabolizable or toxic suppressed the responses of bursting GRNs to sucrose. These abilities to encode information about sugar value are a refinement to the bumblebee’s sense of sweet taste that could be an adaptation that enables precise calculations of the nature and nutritional value of floral nectar.

## Introduction

Sensory acuity evolves in animals when information resolution improves fitness. Visual acuity for colour, for example, arises when the ability to discriminate among colours improves the location of food or the choice among mates (Osorio and Vorobyev, 2008). Where high acuity is necessary to perform optimal stimulus identification, sensory systems become more specialized. While this has been observed in vision, audition, and olfaction, it has relatively rarely been studied in the sense of taste.

The canonical model of the organization of the gustatory system proposes that individual cells or neurons express receptors that detect compounds from specific modalities (Breslin and Spector, 2008; Chen and Dahanukar, 2020). This arrangement permits a form of coding specificity whereby tastants in particular modalities (e.g., sweet, bitter, salty) are represented through the activity of individual neurons. In insects, contact chemosensilla house several (i.e. 2-6) gustatory neurons, each of which responds to compounds from a specific modality (Dethier, 1976). For example, in a fly taste sensillum housing four neurons, one GRN would detect sugars, another ‘low’ salt concentrations, another would detect water, and the last would detect ‘high’ concentrations of salt and/or bitter compounds (Amrein and Thorne, 2005; Chapman and Chapman, 1998; Dethier, 1976). The sensitivity of each neuron type is determined by the receptor proteins (Grs) it expresses.

Sugar compounds are important nutrients for insects. In Drosophila, the GRN in each sensillum that responds to sugars expresses Gr proteins from the 5a, 61a, and the 64a-f family that form complex and diverse heteromeric receptors that enable flies to detect many sugars using the same neuron in each sensillum (Hiroi et al., 2002; Fujii et al., 2015; Dahanukar et al., 2007). These GRNs also express another Gr protein, 43a, which forms functional homomeric receptors tuned towards the detection of fructose (Miyamoto et al., 2012). To date, most studies of sugar detection in insects focus on differences in the tuning of GRNs towards sugars of different structure in order to relate structure to receptor specificity (Dahanukar et al., 2007; Fujii et al., 2015; Miyamoto et al., 2012).

Bees are unique among insects because they possess few genes for gustatory receptors (Robertson and Wanner, 2006; Sadd et al., 2015). Adult bees have specialized mouthparts for collecting floral nectar, a sugary reward offered by plants to pollinators. Sugars have a large impact on bee survival; for example, adult worker honeybees require the greatest amount of ATP of any animal recorded (Suarez et al., 1996). For this reason, one might predict *a priori* that bees would have proportionally greater acuity for sugars in food than other insects, but this has rarely been investigated.

Floral nectar is mainly composed of sucrose, glucose and fructose. However, many bee species collect honeydew produced by Hemipteran insects as a carbohydrate-rich food source and it often contains sugars such as maltose and melezitose (Shaaban et al., 2020). Previously, we found that the bumblebee’s gustatory system exhibits a specialized mechanism for encoding sugar concentration (Miriyala et al., 2018). We observed that two GRNs per sensillum coordinated their activity to produce bursts of spikes at concentrations of sucrose >25 mM. Here, we performed a series of experiments to test the acuity of the gustatory system towards sugars relevant to bumblebees. We measured how compound detection, such as the spiking pattern produced by the sugar-sensing GRNs, was influenced by compound identity. We also tested how each sugar influenced feeding reflexes and bout duration.

## Results & Discussion

### Sugars elicit responses from three of four neurons per galeal sensillum

Bumblebees have a unique mechanism for sensing sucrose in nectar; sucrose evokes a bursting pattern of spikes derived from the activity of two GRNs (GRN1, GRN2) that communicate via a gap junction (Miriyala et al., 2018). This contrasts with the strategy of taste coding described in *Drosophila* where sucrose evokes phasic activity in a single GRN in each sensillum (Hiroi et al., 2004). Temporal codes from peripheral neurons relay important qualitative information about stimuli in other senses including olfaction (Raman et al., 2010). We therefore tested whether temporal coding occurred in bee gustatory neurons. We stimulated bumblebee type A sensilla with seven sugars found in floral nectar and aphid honeydew (fructose, glucose, sucrose, maltose, melezitose, trehalose, sorbitol). We also tested four sugars less commonly encountered by bees (sorbose, xylose, lactose, mannose). Sugars evoked a diverse range of GRN responses that were unrelated to similarity in their compound structure (Figure 1A).

**Figure 1:**
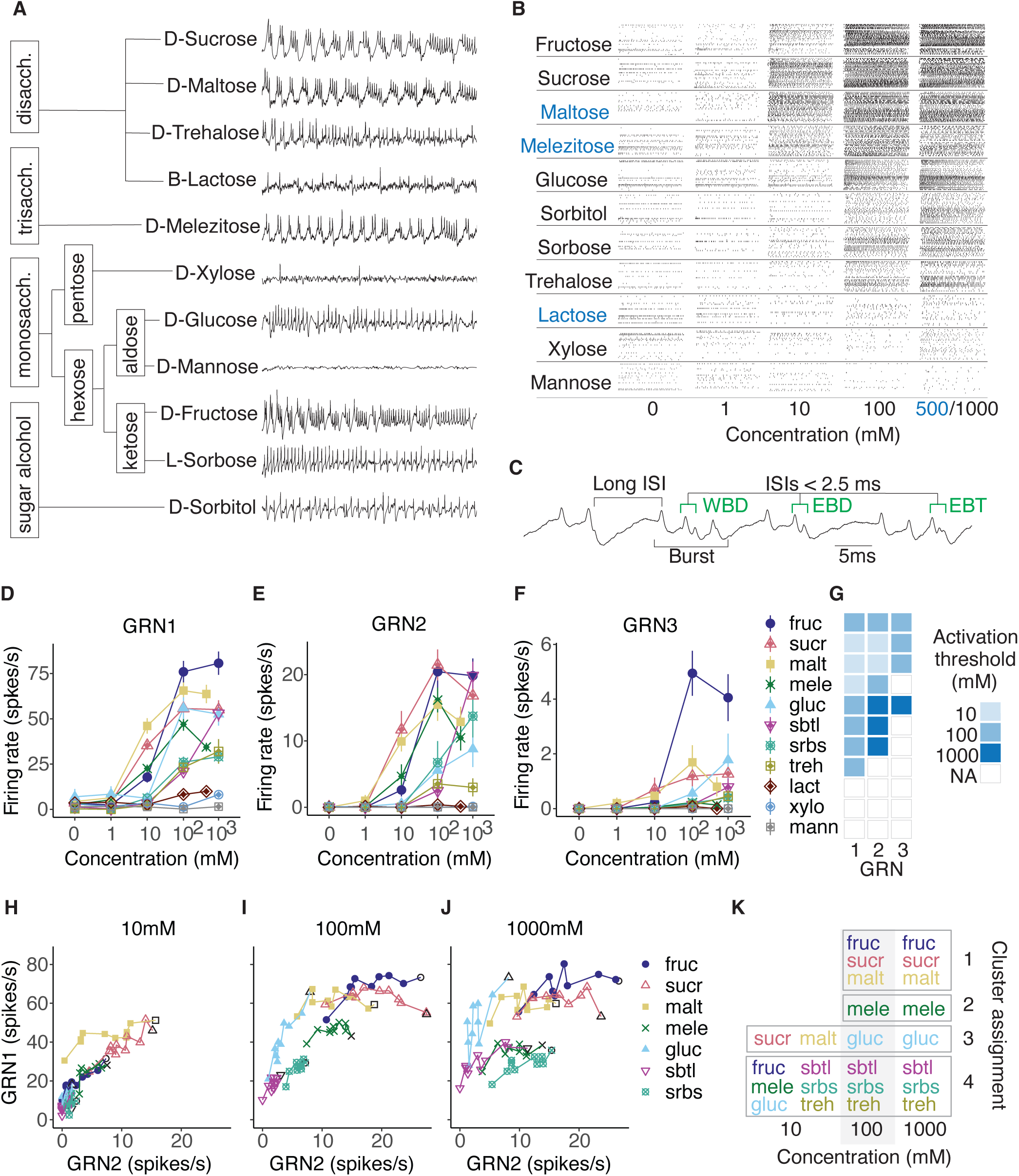
Stimulation with sugars elicits responses from three gustatory receptor neurons per galeal sensillum. **(A)** Sugar structural relationships and representative extracellular tip-recordings of galeal GRNs stimulated with sugars at 100 mM. **(B)** Rasters of GRN spikes over 1 s stimulations with sugars of increasing concentrations from 0 to 1000 mM (Blue lettering is used to indicate sugar stimuli that were 500 mM at their maximum concentration in this experiment due to solubility). **(C)** Spike sorting exploited inter-spike intervals (ISI) to differentiate GRNs: bursts of spikes with 5-10 ms ISIs (GRN1 spikes) were terminated by a single GRN2 spike marked by an end of burst doublet (EBD) with a very short ISI (<2.5 ms) (Miriyala et al., 2018). A third neuron (GRN3) fired occasionally within bursts as a within burst doublet (WBD) or end of burst triplet (EBT) with ISIs <2.5 ms. Bursts were followed by longer (>15 ms) inter-burst intervals. **(D-F)** The average firing rates of GRNs 1-3 over 1 s stimulations varied by sugar type and increased as a function of concentration (n=18 to 24 sensilla from n=6 to 8 bees per sugar). Points represent mean (*±* s.e.m) responses across animals. Concentration response curves for GRNs 1 and 2 varied significantly by sugar treatment (GRN1: F_30,807_=45.55, p<0.0001; GRN2: F_30,837_=23.86, p<0.0001). **(G)** Activation thresholds for GRNs 1-3 for all sugars. Box colours denote significant activation of a single sugar at a given concentration versus water (linear mixed effects, concentration: GRN1: F_4,257_=213.21, p<0.0001; GRN2: F_4,257_=77.98, p<0.0001; GRN3: F_4,257_=20.37, p<0.0001, sugar: GRN1: F_10,66_=23.33, p<0.0001; GRN2: F_10,66_=11.57, p<0.0001; GRN3: F_10,66_=7.58 p<0.0001; conc:sug: GRN1: F_40,257_=13.22, p<0.0001; GRN2: F_40,257_=7.43, p<0.0001; GRN3: F_40,257_=5.48, p<0.0001) with estimated marginal means (EMM) *post hoc* multiple comparisons across concentrations for each sugar (coloured box denotes p<0.05 for a sugar versus water at a given concentration). EMM *post hoc* comparisons between sugars in Figure S1. **(H-J)** Firing rates (spikes per 100 ms bin) of GRN1 versus GRN2 over 1 s stimulations with sugars that significantly activated GRN2 at 10 **(H)**, 100 **(I)**, and 1000 mM (**J**, 500 mM for maltose and melezitose). Points represent mean spikes/s in each bin across all trials for a given sugar, connected with lines over time (over 10 bins). The symbol for the first bin for each sugar (i.e., time = 100 ms) is an open marker with a black outline. GRN2 significantly predicts the trajectory of GRN1 (GLM, GRN2: F_1,194_=1723.0, p<0.0001; sugar: F_6,194_=67.8, p<0.0001; concentration F_2,194_= 138.6, p<0.0001). **(K)** Cluster assignment for each sugar and concentration using the PCs of GRN1 and GRN2 responses in 100 ms bins over 1 s of stimulation. Stimuli were assigned to the cluster to which the largest proportion of individual sensilla responses matched (mean cluster consensus across stimuli: 0.79, CI 0.07). Boxes surround stimuli assigned to a single cluster. Sugars are abbreviated throughout the manuscript as follows: sucr = sucrose, fruc = fructose, malt = maltose, mele = melezitose, gluc = glucose, sbtl = sorbitol, srbs = sorbose, treh = trehalose, lact = lactose, xylo = xylose, mann = mannose.

Sucrose, fructose, and maltose elicited the highest rates of bursting; GRN firing rate also increased as a function of concentration for every sugar tested except lactose, xylose, and mannose (Figure 1B and D-F). We used a spike sorting algorithm that differentiates individual GRNs based on their interspike intervals (ISIs, Figure S1). As expected (Miriyala et al., 2018), we observed coherent bursts of spikes from GRN1 punctuated by a single GRN2 spike. We also detected a third sugar-sensitive GRN in each sensillum (Figure 1C). GRN3 spikes occurred sparsely over the course of sugar stimulation and were not tied to the burst structure of GRN1 and GRN2 (Figure 1C, Figure S1).

Sucrose, maltose and melezitose activated GRN1 and GRN2 at the lowest threshold concentration (10 mM, Figure 1G). Another group of sugars (glucose, sorbitol, and sorbose) activated GRN1 but required sugar concentrations >100 mM to activate GRN2 (e.g., glucose, 1000 mM, Figure 1D-F). The firing rate of GRN3 was low: at the highest concentrations GRN3 fired at 2-6 spikes/s, whereas GRN1 fired at 25-75 spikes/s and GRN2 fired at 4-20 spikes/s (Figure 1D-F). GRN3 responded consistently with a low rate towards fructose but also responded sparsely to sucrose, maltose, and glucose (Figure 1G, Figure S1). None of the GRNs responded to lactose, xylose or mannose.

### Sugars are encoded by a temporal pattern created by bursting neurons

At least three parameters of spike trains encode information: relative spike timing, changes in spike rate (i.e., adaptation) and instantaneous firing rate. GRN responses are typically compared by averaging firing rates over relatively long intervals for neurons (e.g, 1 s, Figure 1D-F). Bumblebee GRNs, however, clearly have a temporal pattern of firing which could contribute to coding over shorter time scales (Figure 1A). Sugar-induced changes to the temporal structure of GRN firing is readily visualized in the traces in Figure 1A.

Analysis of the relationship between the firing of GRN1 and GRN2 over the time course of sugar stimulation revealed that concentration and sugar identity affect the bursting pattern of activity (Figure 1H-J). GRN1 spike rate increased as a function of GRN2 for all 10 mM stimuli (except maltose) with an average slope of 2 (Figure 1H and Table S1). At >100 mM (Figure 1I and 1J), sugars which elicited regular and relatively high rates of GRN2 spikes (fructose, sucrose, maltose, melezitose) became distinguishable from all other sugars because the rate of GRN1 depends less on the rate of GRN2 firing. Finding that the dynamic response between GRN1 and GRN2 depends on sugar identity (as in Figure 1J) and not entirely on the rate of GRN1 firing indicates that GRN2 also plays an active role in sensing sugars.

The temporal activity of GRNs could potentially reveal information about a sugar’s chemical identity (Reiter et al., 2015) (Figure S1). To test this, we employed a consensus clustering analysis (CCA) of output from a principal components analysis of the binned spike trains of GRNs 1 and 2 (as in Stopfer et al (Stopfer et al., 2003), Figure S2 and Table S2). Sugars that did not evoke spikes such as xylose, mannose, and lactose were not included in the analyses. The CCA predicted how the pattern of spiking, rates of spiking, and involvement of all three neurons encoded information about sugar stimuli (Figure 1K). At 10 mM, the CCA was only able to identify two groups: one for maltose and sucrose and one that contained all the other sugars (Figure 1K). At concentrations >10 mM, the CCA identified four distinct groups. Cluster 1 was composed of fructose, sucrose, and maltose. Responses to these sugars exhibited high rates of burst firing in GRN1 and GRN2. Cluster 2 represented melezitose, which evoked high rates of burst firing, but fewer GRN1 spikes per burst (lower GRN1 rate). (Note: sugars in Clusters 1 and 2 were also more likely to evoke GRN1 spikes at 10-100 mM, Figure 1G). Cluster 3 represented glucose. Glucose evoked relatively high firing activity in GRN1, but less activity in GRN2. Furthermore, glucose responses also exhibited a strong rate of adaptation, as indicated by the slope of the relationship between GRN1 and GRN2 in Figure 1D-F (Table S1). Cluster 4 represented trehalose, sorbitol, and sorbose. Although sorbitol and sorbose elicited high GRN2 firing rates at 1000 mM, this did not correspond with high rates in GRN1 (Figure 1J). The clustering of sugar-evoked responses revealed that the pattern of spiking was an important facet of the code for sugars.

### Nutritive value of sugars predicts palatability

Taste instructs feeding behaviour. To identify how input from the mouthparts influenced the relative phagostimulatory power of each sugar, we used two additional behavioural assays. Bees reflexively extend the proboscis to feed when the sensilla on the mouthparts are stimulated (Miriyala et al., 2018). All of the 500/1000 mM sugar stimuli that elicited GRN spikes (Figure 1) also provoked proboscis extension when the mouthpart sensilla were directly stimulated, with the exception of sorbitol (Figure 2C). Furthermore, the phagostimulatory properties of the sugars also affects the duration of feeding once it commences (Ma et al., 2016). We used a 2 min feeding assay to assess how the structure of feeding was affected by the sugars. At 100 mM, the only sugars that bees consumed in significant volumes were fructose, sucrose, maltose and melezitose, however bees only maintained contact with sucrose and maltose solutions significantly longer than water (Figure 2D). Bees consumed >10 µl of all of the 500/1000 mM sugar stimuli except lactose, xylose, sorbitol and mannose (Figure 2E). They also spent more time in contact with the solution and had longer first bouts on the sugars that provoked the highest spike rates in the GRNs (Figure 2E).

**Figure 2:**
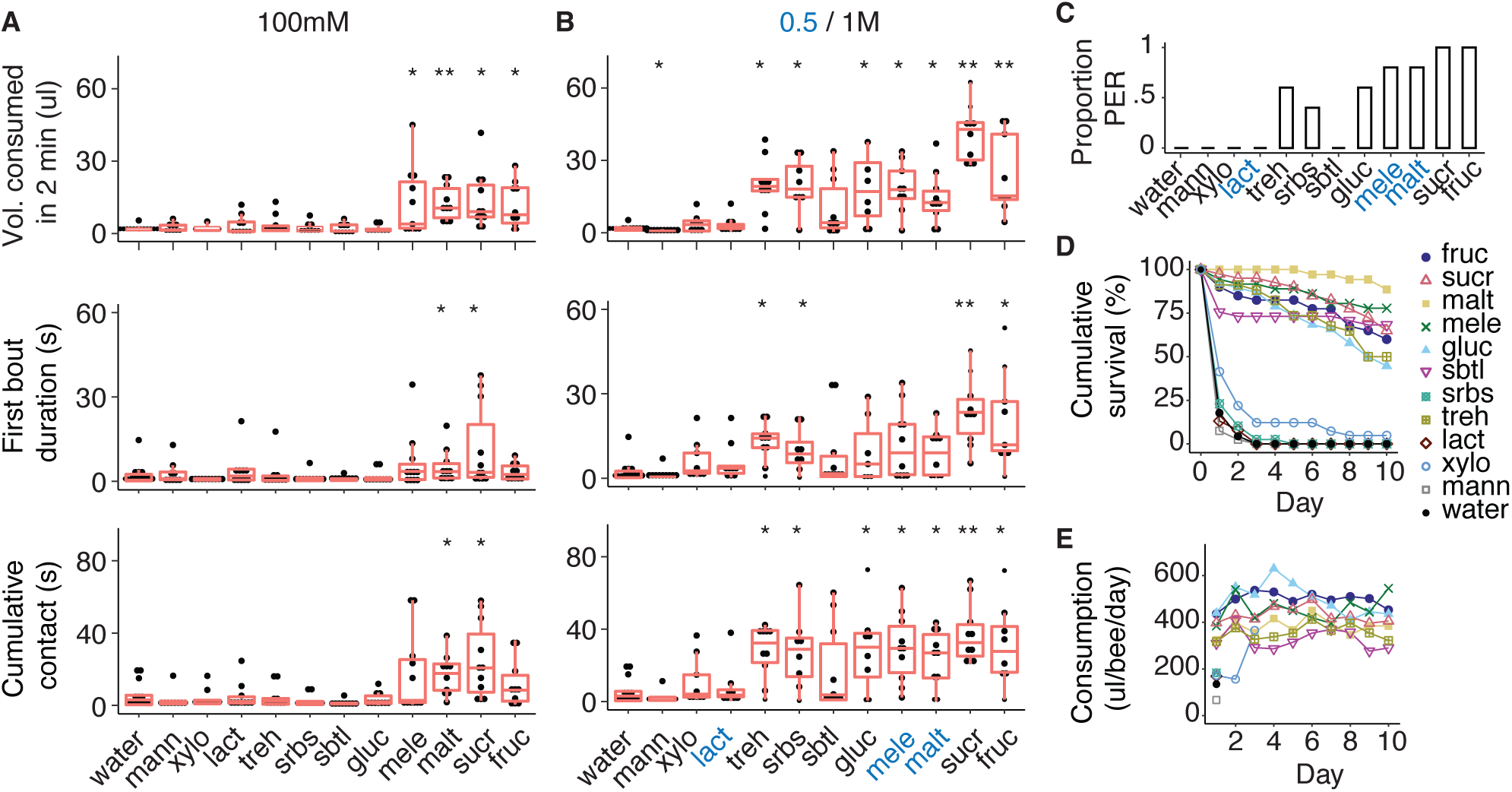
Nutritional value of sugars corresponds with palatability. **(A-B)** Feeding behaviour over 2 min varies by sugar type. Bees consumed more of the sugar solutions at the highest concentrations (top, 100 mM, **A**, Kruskal-Wallis, 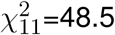, p<0.0001, n=10 to 13; 1000 mM, 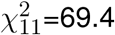, p< 0.0001, n=8 to 13 per sugar), and spent more time in contact with the most phagostimulatory sugars over the first bout (middle, 100 mM, 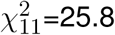, p<0.05, n=10 to 13; 1000 mM, 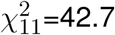, p<0.0001, n=8 to 13 per sugar) and entire 2 min period (bottom, 100 mM, 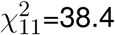, p<0.0001, n=10 to 13; 1000 mM, 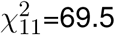, p<0.0001, n=8 to 13 per sugar). Blue letters denote 500 mM concentrations. Asterisks represent significant differences in sugar consumption versus water (Wilcoxon rank-sum multiple comparisons). **(C)** The probability of eliciting the proboscis extension reflex depended on sugar type (Kruskal-Wallis, 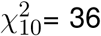, p<0.0001, n=5 per sugar). **(D)** Survival of bumblebees fed with 1000 mM (or 500 mM for lactose, maltose, and melezitose) solutions depended on sugar type (Kaplan-Meier Log-Rank statistic =1576, df=11, p<0.0001). Holm-Sidak *post hoc* analysis showed significant differences in survival between the following groups of sugars (from highest to lowest survival): 1) maltose; 2) fructose, sucrose, melezitose; 3) glucose, sorbitol, trehalose; and 4) sorbose, lactose, xylose, mannose and water. (n=33 to 42 bees per treatment). **(E)** Total consumption of sugar solutions per 24 h period over the 10-day longevity assay in D.

Whether or not a bee feeds on a solution is likely to be related to the metabolic or nutritional value of a sugar. We fed bees with 500/1000 mM solutions of each of the sugars and measured their lifespan over a 10-day period. At least 60-80% of the bees fed with fructose, sucrose, melezitose, or sorbitol, survived for 10 days, whereas nearly 90% of the bees fed with maltose survived (Figure 2F). Unexpectedly, only 50% of the bees fed with glucose or trehalose survived over the 10 day period. In contrast, most bees fed with sorbose, lactose, xylose or mannose died within 3 days of the start of the experiment (Figure 2F). These bees did not consume much of the solution and died of starvation or of toxicity (Figure 2G).

### GRN3 is not activated by ligands from other modalities

Our data indicate that three of the four GRNs in bumblebee galeal sensilla respond to sugars. This finding contrasts the classical model in which insects possess one sugar-sensing GRN per sensillum (Thorne et al., 2004). To our knowledge, the only other example of an insect with multiple within-sensillum GRNs sensitive to sugars is in hawkmoth larvae (*Manduca sexta*). The taste sensilla of these caterpillars contain at least two GRNs that are responsive to sugars. However, these GRNs do not burst, and dual activation of GRNs occurs primarily with sugar mixtures rather than single sugars (Glendinning et al., 2007). GRN1 and GRN2 clearly demonstrate modulation of firing activity consistent with sugar-sensing. However, the low rate of GRN3 firing to sugars might indicate that it responds more strongly to other specific ligands. The tip-recording technique does not always obtain/retrieve traces that permit the identification of GRNs 1-3 responses by the shape of their spikes. For this reason, we could not determine whether GRN3 responded to other ligands presented as monomolecular stimuli. We instead tested the responses of the GRNs to fructose mixed with salts (NaCl, KCl), bitter compounds (caffeine, quinine), or amino acids (glutamic acid, proline, phenylalanine) to identify if GRN3 could be activated by ligands from other taste modalities (Figure 3A). As shown in other studies in bees, quinine completely silenced all GRN responses to fructose (Wright et al., 2010; Kessler et al., 2015). None of the other compounds influenced GRN3 firing activity (Figure 3B). The pattern of activity of GRN1 and 2 over time was moderately affected by proline (Figure S3), which reduced GRN2 firing rates and affected bursting structure by reducing the number of GRN1 spikes per burst and increasing its rate of adaptation (Figure S3). However, when the patterns of activity of all three GRNs were integrated into a CCA of the binned spikes, none of the stimuli were significantly grouped into a distinct cluster except for the fructose-quinine mixture (Figure 3C-D, Figure S3).

**Figure 3:**
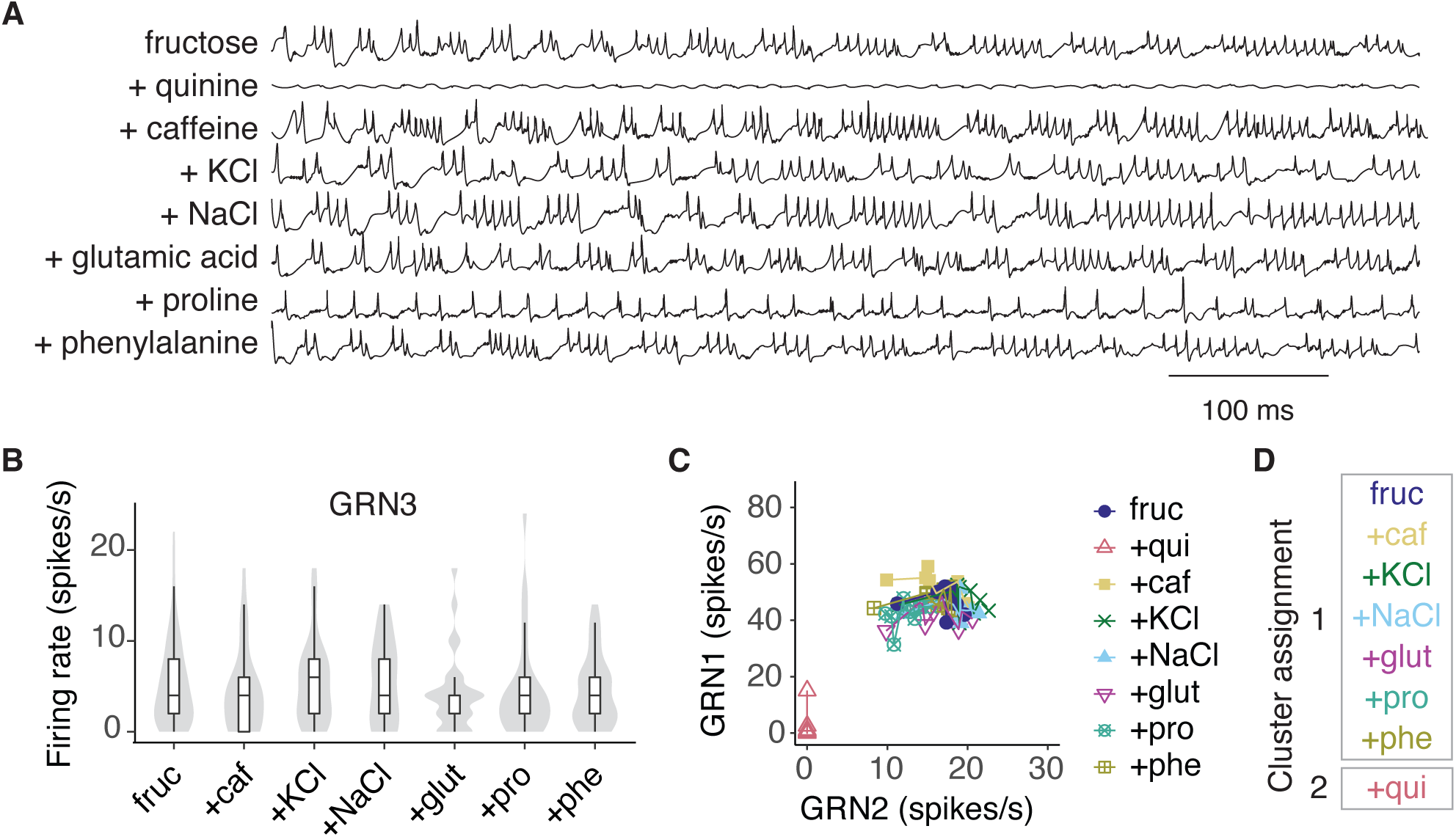
GRN3 responses are not affected by the addition of secondary compounds in fructose. **(A)** Representative tip-recordings from galeal A-type sensilla stimulated with 100 mM fructose, or 100 mM fructose plus 1 mM bitter compound (quinine, caffeine), salt (KCl, NaCl), or amino acid (glutamate, proline, phenylalanine). Individual sensilla were tested with all compounds (in total 24 sensilla, n=8 bees). **(B)** Average GRN3 spike rates (over 1 s stimulation) to fructose are not affected by the addition of secondary compounds (linear mixed effects, F_6,267_=0.452, p>0.05). Responses to quinine not included; quinine shuts down all GRN responses. **(C)** Firing rates (spikes per 100 ms bin) of GRN1 versus GRN2 over 1 s of stimulation with fructose and fructose mixtures. Points represent mean spikes/s in each bin across all trials for a given sugar, connected with lines over time (over 10 bins). GRN2 significantly predicts the trajectory of GRN1 over time (GLM, GRN2: F_1,65_=786.1, p<0.0001; sugar F_6,65_=36.0, p<0.0001). **(D)** Responses to fructose and fructose mixtures cluster in two groups (with or without quinine) using a consensus clustering analysis of the binned GRN responses (Figure S3).

### Non-nutritive sugars reduce GRN responses to sucrose

In the previous experiments, we established that sugars with nutritional value activate GRNs 1-3. Some of the sugars we tested did not activate any GRNs. To test whether activity evoked in GRNs 1-3 relates to the nutritional value or toxicity of the sugars, we fed bees with sorbose, xylose, lactose, or mannose in equimolar mixtures of the sugars with sucrose. Xylose had some nutritional value, as bees fed with sucrose mixed with xylose lived longer than those fed with sucrose alone. Sorbose provided no additional nutritional value, and it was not toxic. However, bees fed with sucrose laced with mannose and lactose died at a much faster rate, indicating that these sugars were toxic (Figure 4A). In a 2 min assay of feeding, the addition of mannose, lactose and xylose to sucrose did not significantly alter the quantity of food a bee would consume (Figure 4B). However, bees fed sucrose containing mannose over a 24 h period ate <30% of the total volume consumed by bees fed with the other sugar mixtures (Figure 4C) indicating that post-ingestive feedback from the toxicity of mannose inhibited ingestion. Unlike sorbose, the sugars mannose, xylose, and lactose did not evoke feeding or spikes, but might still be detectable in sugar solutions. To test this, we stimulated the sensilla with mixtures of sucrose and each individual sugar. As a positive control, we also tested a fructose-sucrose mixture. The firing rates of GRN1 and GRN2 to sucrose were attenuated by the addition of mannose, lactose, and xylose. Sorbose reduced the rate of GRN1 firing but not the rate of GRN2 firing. In contrast, the addition of fructose increased the rate of firing of both GRN1 and GRN2 (Figure 4D). Although addition of the sugars to sucrose affected GRN1 and GRN2 average firing rates, the pattern of activation of these GRNs was similar across stimuli (Figure 4E, Figure S4). The CCA did not distinguish responses of sensilla to the various sugar mixtures (Figure 4F, Figure S4). This clustering predicts that, although the mixtures have effects on absolute GRN firing rates, bees cannot distinguish between these mixtures based on their taste alone.

**Figure 4:**
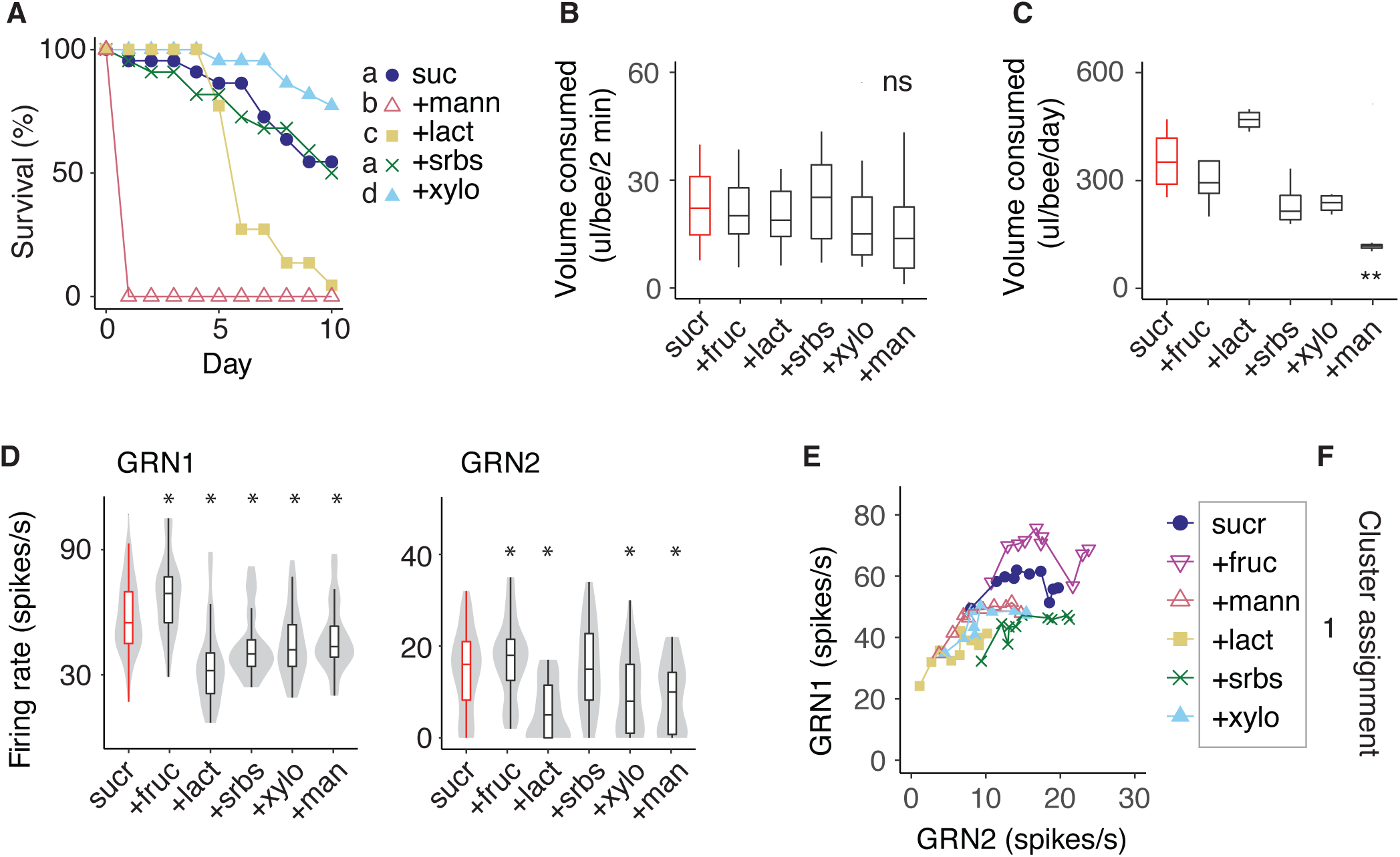
GRN responses to sucrose are attenuated by the addition of non-nutritive secondary compounds. **(A)** Bees fed with sucrose mixed with a non-nutritive sugar (500 mM equimolar solutions) over 10 days did not survive if the mixture included mannose and lactose, but did not have a different rate of mortality if mixed with sorbose. Those fed with sucrose mixed wtih xylose were more likely to survive than the control. Letters denote significant differences in survival (Kaplan-Meier log-rank statistic =370.9, df=4, p<0.001, n=20 to 23 bees per sugar). **(B)** Feeding behaviour over 2 min did not vary in the total volume consumed with the addition of a non-nutritive sugar (ANOVA: F_5,88_=1.66, p=0.15, n=15 to 17 per sugar solution). **(C)** Over 24 h, the quantity of food consumed per bumblebee was significantly less if the bees had been fed with the sucrose-mannose mixture (ANOVA with Tukey multiple comparisons, F_5,18_=10.53, p<0.001, n=4 cages, 20 bees per treatment). Asterisks represent significance versus sucrose alone. **(D)** Average GRN1 (left) and GRN2 (right) firing rates following stimulation with sucrose (500 mM) and equimolar (500 mM) sugar mixtures, including a sucrose-fructose mixture (Aligned Rank Transform, GRN1: F_5,263_=26.8, p<0.0001; GRN2: F_5,258_=19.7, p<0.0001). Violin plots show the distribution of responses of individual sensilla, with boxplots overlaid. Asterisks denote significant differences of GRN responses to the mixture versus sucrose (red boxes). **(E)** Firing rates (spikes per 100 ms bin) of GRN1 versus GRN2 over 1s of stimulation with sucrose and sucrose mixtures. Points represent mean spikes/s in each bin across all trials for a given sugar, connected with lines over time (over 10 bins). GRN2 significantly predicts the trajectory of GRN1 over time, slopes do not vary significantly between sugars (GLM, GRN2: F_1,48_=195.7, p<0.0001; sugar F_5,48_=36.8, p<0.0001, GRN2*sugar F_5,48_=214.9, p>0.05). **(F)** The consensus clustering analysis of the binned GRN responses to sucrose and sucrose mixtures did not partition the stimuli into separate groups.

## Conclusions

Our data here indicate that the temporal pattern of firing of GRNs in the bumblebee’s galeal sensilla has the capacity to partition sugars into at least 4 categories according to their nutritive value: ‘very valuable’ (fructose, sucrose, maltose), valuable (melezitose), slightly less valuable (glucose), or not very valuable (trehalose, sorbose, sorbitol). In addition, the galeal GRNs do not respond to toxic or low value sugars (xylose, mannose and lactose). The bee’s mouthparts have hundreds of taste sensilla in other locations, such as the labial palps and glossa. Sensilla in other locations on the mouthparts likely contribute to the resolution of sugars in the brain, consistent with the analysis of taste input from the maxillary nerve of adult hawkmoths by Reiter et al (Reiter et al., 2015). Our data suggest bumblebees may be capable of distinguishing sugars by identity, although this remains to be tested.

Fine tuning of the gustatory system towards high-value sugars is likely to provide bees the ability to select the optimally formulated nectar to fuel flight. This is particularly important for a small flying animal to derive the maximum energy from the least weight that it can manage. Interestingly, although fructose and glucose have the same molecular weight and are both found in nectar, fructose elicits higher GRN responses and is more phagostimulatory. Unlike fructose, glucose crystallizes at low temperatures, making it harder to drink from cold nectar solutions (Silva et al., 2010). Fructose is also particularly valuable to bumblebees because they use it as the substrate for activation of flight muscles to heat up during ‘shivering’ (Beenakkers et al., 1984). The only sugar for which we did not see a relationship between GRN activation and survival was sorbose. Sorbose elicited PER and drinking behaviour in the first bout of feeding but it was not consumed in large volumes over 24 h (Figure 2G). For this reason, bees fed sorbose did not live longer than 4 days (Figure 2E). Likewise, bees that consumed mannose mixed with sucrose stopped feeding only after they had consumed the solution. These data indicate that post-ingestive feedback plays an important role in modulating feeding towards sugars over periods as long as 24 h. Integration of taste information with post-ingestive cues, therefore, is likely to improve the accuracy of the brain to identify the nutritional value of foods which are often mixtures of substances.

## Methods

### Resource availability

#### Lead contact

Further information and requests regarding the methods used should be directed to the lead contact, Geraldine A. Wright.

#### Materials availability

This study did not generate new unique reagents.

#### Data and code availability

All (.csv) data files have been deposited at *[to be confirmed]* and are publicly available as of the date of publication. Accession numbers are listed in the key resources table. The original code for consensus clustering has been deposited at *[to be confirmed]* and is publicly available as of the date of publication. DOIs are listed in the key resources table. Any additional information required to reanalyze the data reported in this paper is available from the lead contact upon request.

### Experimental Model and Subject Details

#### Bumblebee colonies

Bumblebees Experiments were performed at Newcastle University and the University of Oxford with commercial colonies of *Bombus terrestris audax* (Koppert Biological Systems, Haverill, UK and Biobest, Westerlo, Belgium). Bumblebees were kept inside their colonies and maintained at laboratory conditions (22-27°C and 25-40% RH) until use for experiments. They were fed ad libitum with honeybee-collected pollen (Koppert Biological Systems, Haverill, UK and Agralan, Swindon, UK) and the proprietary sugar syrup provided with the colonies (Biogluc, Biobest) or 1 M sucrose. Workers were collected directly from the colony. To minimize the likelihood of nurse bee inclusion, only bees with a thorax width >4.5 mm were used in experimentation (Goulson et al., 2002). Bees used were of unknown age and were experimentally naïve. Treatments were counterbalanced across colonies and bees from within a colony were randomly allocated to a treatment group for all experiments. Health status of the colonies was unknown but they are presumed to be disease-free due to the strict quality controls conducted by commercial bumblebee suppliers (Huang et al., 2015).

## Method Details

### Sugar Solutions

Solutions of sugars were prepared in demineralised water for both electrophysiological and behavioural experiments. Sugars were purchased from Sigma-Aldrich (Dorset, England) or Alfa Aeasar (Heysham, UK) at a 98% minimum purity. One trisaccharide [D-(+)-melezitose monohydrate], four disaccharides [D-(+)-sucrose, D-(+)-maltose monohydrate, D-(+)-trehalose, β-lactose], five monosaccharides [D-(-)-fructose, D-(+)-glucose, D-(+)-mannose, L-(-)-sorbose, D-(+)-xylose] and one sugar alcohol [D-sorbitol] were tested.

### Proboscis extension response

To test the phagostimulatory effect of sugars, proboscis extension response (PER) experiments were performed on bumblebees. Individual bees were collected directly from the colony in small glass vials and cold anaesthetized on ice for approximately 3 min, or until movement slowed sufficiently, before being harnessed. Harnesses were modified 1000 µl pipette tips with a portion of the tip removed to allow placement of the bee. A single duct tape strip (2 mm in diameter) was placed across the bees thorax to hold the animal in place. A small piece of wire was positioned directly below the bees mouthparts and was affixed to the harness with melted beeswax. The wire functioned to stop the bee from folding the mouthparts below their mandibles. Following harnessing, bees were starved for 3-8 h at room temperature in a dark environment to habituate them to the harnesses and to motivate them to feed during the assay. A 1.0 ml syringe with a female adapter was used to deliver a small drop (3.5 µl) of solution directly to the bees mouthparts for a period of 4 s and the PER was monitored visually by the same experimenter. The test solution was tested first, followed by a water control and a 1000 mM sucrose control. Five bees were tested for each sugar tastant. A PER was counted if the bee extended its proboscis to at least 90° with the glossa passing the point between the mandibles. Bees that displayed extensive licking behavior of the glossa immediately before or during the assay were excluded as this behavior can confound accurate PER quantification. Animals that did not display the PER to 1000 mM sucrose were excluded from analyses, as were those that displayed the PER to water. Experimenters were not blind to the treatments.

### Capillary taste assay

The feeding behaviour of free-moving bumblebees was examined in more detail using a capillary taste assay previously described in Ma et al (Ma et al., 2016). Individual bees were collected directly from colonies using small plastic vials (7 cm long, 2.8 cm inner diameter, with a perforated plastic stopper) and placed in a dark box maintained at laboratory conditions (24°C and 38% RH) for 3-6 h to starve them sufficiently to ensure their motivation to feed during the assay. Following starvation, bees were transferred into a modified 15 ml centrifuge tube (119 mm length, 17 mm diameter), with a 3 mm hole drilled at the tip and a steel mesh (10*×*30 mm, 1 mm mesh diameter) inserted at the bottom of the tube to allow the bumblebees to grip to the wall of the tube. Bumblebees were free to move within the holding tubes throughout the experiment. The tube was then placed into a polystyrene holder and held in place using dental wax. White cardboard affixed to either side of the holder shielded the bees from visual stimuli. Bees were allowed a 3 min habituation period to the test environment before being presented with a droplet of 500 mM sucrose (3.5 µl) using a 1.0 ml syringe connected to a female adapter. The droplet served to bait the bees to extend their proboscis. If the droplet was not consumed within 5 min, the bees were excluded from the experiment. Immediately after consuming the sucrose droplet the bees were offered a 100 µl microcapillary tube 60 mm in length (Blaubrand intraEnd, Ref 7091 44) filled with the test solution.

Solutions were blinded for this assay. The microcapillary tube was held in place by feeding the tube through a modified 1.0 ml syringe with the tip removed. The syringe was in turn affixed to a micromanipulator using dental wax. The proximity of the solutions to the bees’ mouthparts could be controlled by adjusting the micromanipulator and by squeezing a silicone tube (60 mm long, 1 mm inside diameter) which allowed the microcapillary tube to function as a pipette bulb. The microcapillary was adjusted such that the solution was always readily accessible to the bee. The feeding period lasted for 2 min after the bumblebee tasted the solution within the microcapillary with its mouthparts. Bumblebees failing to taste the test solution with their mouthparts within 10 min of being presented the test solution were discarded from the experiment. The movement of the mouthparts was recorded using a digital microscope (Dino-Lite AM4815ZT) which was positioned 20 cm above the end of the microcapillary tube. Video recordings were made using DinoLite Digital Microscope software at resolution of 640×480 pixels, 26 Hz frame rate, and 25x magnification. Videos were manually scored at 0.5x speed for contact times of the mouthparts with the feeding solution using the Noldus Observer software (The Observer® XT 5.0.25, Noldus, Wagenigen, The Netherlands). A feeding bout was defined as a contact between the extended proboscis and the solution not interrupted by an absence of contacts of 5 s or more. For each sugar and concentration n=9 to 12 bees were tested.

Each microcapillary tube was scanned at 600 d.p.i. before and after the experiment, and fluid levels were measured using ImageJ (version 1.48) (Schneider et al., 2012). The reference scale was set to 60 mm. Image files were zoomed to approximately 400% and the length of the solution was measured meniscus to meniscus. The length of test solution consumed by each bee was calculated as the difference between the length of the liquid before and after the test phase. These lengths were converted to volumes using the formula:

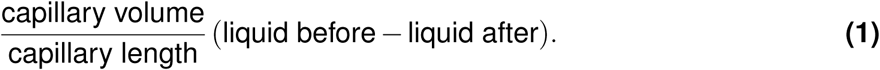

Evaporation was monitored by measuring liquid levels in 34 capillaries filled with water fixed on the experimental set up for a 12 min period (the maximum duration of an experimental trial) in the absence of a bee. Evaporation value appeared to be marginal (0.9*±*0.3 µl, mean *±* s.e.m.). Water consumption was consequently marginal, but slightly higher than evaporation (1.7*±*0.4 µl).

### Longevity assay

The ability of sugars to support bumblebees’ survival was tested. Bees were collected and caged in an identical fashion to that of the 24-hour preference assay, with the exception that bees were constrained to a single diet for the duration of the experiment. The experiments ran for a total of 10 days. Mortality was assessed every 24*±*2 h from the beginning of the experiment and dead bees were removed from the cage using forceps. A piece of paper was placed in the bottom of each cage and replaced when needed to maintain hygienic conditions. Feeding tubes were weighed and replaced every 24 h to monitor the total volume of solutions consumed over the course of the experiment, as detailed earlier. Longevity of bumblebees with access to 500 mM melezitose or lactose, or 1000 mM sucrose, trehalose, maltose, fructose, glucose, xylose, mannose, sorbose, sorbitol, or water were measured, and n=33 to 42 bees were tested for each sugar. In the toxicity experiment, longevity of bumblebees having access to equimolar mixtures (500 mM) of sucrose and lactose, sucrose and mannose, sucrose and sorbose, sucrose and xylose and 500 mM sucrose alone was assessed during 10 consecutive days (n=20 bees per sugar mixture). Experimenters were not blind to the treatments.

### Electrophysiology

To investigate how sugar perception is encoded by GRNs on the mouthparts, tip recordings Hodgson et al. (1955) were performed on individual galeal sensilla chaetica A-type contact chemoreceptors (Supplemental Figure 1A>). Electrophysiological preparations were made as already described (Miriyala et al., 2018; Kessler et al., 2015). For tip recordings, sensilla were stimulated using a motorized micro-manipulator (MPC-200, Sutter Instruments, USA) for 2 s with a borosilicate (Clark capillary glass 30-066, GC150TF-10) recording electrode (15 µm tip diameter, made with a Narishige PC-10 electrode puller) containing the test solution. Signals were acquired using a non-blocking pre-amplifier (TasteProbe; Syntech, Germany), amplified (AC amplifier 1800, A-M Systems, USA), digitized at 10 or 30 kHz (DT9803 Data Translation) and stored using dbWave (version 4.2014.3.22) or DataView (version 11.5). A minimum latency period of 3 minutes was allowed between simultaneous recordings to avoid adaptation. Recordings were made from among the first 6 most distal sensilla chaetica on the galea of bumblebees. For each sugar, n = 6 to 8 bumblebees were tested.

### Spike detection

Both spikes and bursts were detected using custom routines written in Matlab (The MathWorks) following a modified analysis described in Miriyala et al (Miriyala et al., 2018). The first 2 s following the contact artefact (automatically detected with findpeaks [fixed thresholds: minpeakheight = 1.5, npeaks = 1]) were first band-pass filtered at 300-2500 Hz (second order butterworth filter) with the butter function. Assuming a normal distribution of noise frequencies, the standard deviation of the background noise was estimated using:

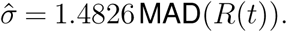

Each recording was then normalised using:

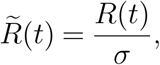

where *R*(*t*) is the filtered recording and MAD the median absolute deviation. Spikes were then detected using the peakfinder function with adequate thresholds. A logarithmic equation of the form:

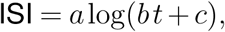

where ISI are the interspike-intervals and *t* are the spike time indices, was used to detect the end of burst positions (EOB). A spike was considered as EOB if its ISI exceeded two times its fitted value. The number of spike doublets (2 spikes events separated by an ISI <2.5 ms) were quantified within each burst (i.e., after removing the last spike event of each burst). The total number of spikes, bursts and doublets within each burst were quantified during a 0.1 to 1.1 s recording window following the peak of the contact artefact. GRN1 spikes were identified as any low-frequency responses to sugars, and as the high frequency component of the bursts, while GRN2 spikes were identified exclusively at the EOB position, as described in Miriyala et al (Miriyala et al., 2018). The presence of within burst doublets and end of burst triplets were identified in the responses to some sugars, representing the spikes from GRN3.

### Spike train analyses

We compared the average firing rates of GRNs housed in sensilla 1-6 in response to water and found that sensilla 4-6 responded with higher firing rates than sensilla 1-3 (Supplemental Figure 1B). Sugars that did not elicit strong GRN1 responses (lactose, xylose, mannose) shut down the water cell at concentrations above 10 mM (Supplemental Figure 1C). Spike sorting methods could not differentiate the GRNs responding to water versus GRN1 spikes, so the water response resulted either from activity in GRN1, or another GRN with a similar amplitude, and this presented a confound in measuring sugar responses. Sensilla 4-6 were removed from analyses to define the “activation threshold” (lowest concentration that elicited a significant GRN response compared to water). Responses of GRNs 1-3 from sensilla 1-3 were compared using the number of spikes from 0.1 to 1.1 s from stimulus onset, i.e., the average firing rate. To assess the importance of time-dependent GRN responses, we additionally compared the spike trains of GRN1 and GRN2, by binning spikes for each 100 ms of recording from 0.1 to 1.1 s after stimulus onset. Time series of GRN spikes in 100 ms bins were aligned so that a single sensillum response was contained in a single vector (i.e., 20 features per vector containing binned normalized spike counts for GRN1 and GRN2).

### Quantification and Statistical Analysis

#### Consensus clustering

For subsequent analyses, we focused on the responses to sugars that elicited significant GRN responses only (trehalose, sorbitol, sorbose, glucose, melezitose, sucrose, maltose and fructose at concentrations ≥10 mM). Dimensionality of the GRN time series responses (normalized to peak GRN firing rates) was reduced with a principal component analysis. We used a consensus clustering analysis (CCA) to assess whether GRN responses to sugars were differentiable based on sugar identity or intensity. CCA is an unsupervised class discovery algorithm for determining cluster count and membership by aggregating the results from many clustering iterations to detect possible groupings of items based on intrinsic features. We used the `ConsensusClusterPlus` package in R (Wilkerson and Hayes, 2010), to implement consensus clustering. We used the first eight principal components from the GRN time series data for clustering analyses. We clustered the GRN responses to the sugars that elicited significant GRN responses. We selected a k-means clustering algorithm and ran the consensus clustering algorithm with 2000 iterations, k=2 to 8 clusters, and 90% of the samples and features per iteration (i.e., 10% of the responses and 10% of the bins were omitted from each clustering run). Optimal cluster count (k) was determined using the consensus matrix, cumulative distribution function, as well as cluster and item consensus. The stimuli were grouped by assessing the individual cluster assignments for the responses of individual sensilla: variability in responses between sensilla resulted in some scattering of the responses across different clusters. The stimulus was assigned to the cluster that represented the greatest proportion of the responses across sensilla. We performed a sensitivity analysis by adjusting the hyperparameters of the clustering analysis (i.e., by varying the proportion of features and samples included in each iteration between 0.8 and 1) and additionally by running the clustering analysis on the normalized binned spikes (rather than the principal components), with similar results. Consensus clustering was also used to compare the time series responses of GRNs stimulated with fructose and fructose mixtures, as well as sucrose and sucrose mixtures. These analyses were performed as described above.

#### Other statistical analyses

Statistical analyses were performed in R (version 4.0.3). All data were tested for normality and equal variance using the Shapiro-Wilks and Levene’s tests. Parametric repeated measures data, including average firing rates of GRNs 1-3, maximum number of spikes per burst, and GRN adaptation rates, were modeled using linear mixed effects models (lmerTest package, (Kuznetsova et al., 2017)), with animal ID as a random effect. Estimated marginal means (emmeans package, (Lenth et al., 2018)) was used for *post hoc* multiple comparisons of linear mixed models. The aligned rank transform (ARTool package, (Wobbrock et al., 2011)) was used to transform non-parametric data, including the water response of sensilla 4-8, and fructose mixtures electrophysiological responses, for modeling in a linear mixed model with animal ID as a random effect. Dose response curves for (Figure 1D-E) were modeled with and without sugar as a variable, and these models were compared to quantify differences in the average GRN responses for each sugar (drc package, (Ritz et al., 2015)). The Kruskal-Wallis test (with Wilcoxon Rank Sum *post hoc* multiple comparisons) was used to compare responses of sensilla 1-6 to water, and compare 24 h feeding preferences and consumption of sugars over 2 min. The Kaplan Meier Log Ranks test was used to assess differences in survival of bees fed different sugars or sucrose mixtures. Wilcoxon signed-rank tests were used to test if the median preference index for each sugar pair was significantly different from zero. Where appropriate, p-values were adjusted for multiple testing using Bonferroni corrections within each assay. Significance was assessed at α<0.05. The results of *post hoc* analyses are denoted either by letters to show significant differences between groups, or with asterisks: ‘*’ <0.05; ‘**’ <0.001; ‘***’ <0.0001.

## Acknowledgements

The authors thank Max Stanyard and Mushtaq Al-Esawy for help with pilot experiments for the nutritional value of sugars. This research was supported by a Leverhulme Trust (RPG-2012-708) and two BBSRC grants (BB/M00709X/1 and BB/S000402/1) to GA Wright, and an NSERC PDF (PDF-546125-2020) and Royal Society Newton Fellowship (NIF/R1/19368) to RH Parkinson.

## Author contributions

RH Parkinson, SC Kessler, A Miriyala, and GA Wright conceptualized and designed the experiments. RH Parkinson, SC Kessler, J Scott, J Bu, and A Simpson collected data. A Mahdi assisted in implementing the consensus clustering method. RH Parkinson performed all other analyses and prepared all visualizations. RH Parkinson and GA Wright wrote the manuscript. All authors contributed to revising the manuscript.

